# Benchmarking Large Language Models for Predictive Modeling in Biomedical Research With a Focus on Reproductive Health

**DOI:** 10.1101/2025.07.07.663529

**Authors:** Reuben Sarwal, Victor Tarca, Claire Dubin, Nikolas Kalavros, Gaurav Bhatti, Sanchita Bhattacharya, Atul Butte, Roberto Romero, Gustavo Stolovitzky, Tomiko T. Oskotsky, Adi L. Tarca, Marina Sirota

**Author notes:** co-first author (Reuben Sarwal, Victor Tarca). co-senior author (Adi L. Tarca, Marina Sirota).

## Abstract

Generative AI, particularly large language models (LLMs), is increasingly being used in computational biology to support code generation for data analysis. In this study, we evaluated the ability of LLMs to generate functional R and Python code for predictive modeling tasks, leveraging standardized molecular datasets from several recent DREAM (Dialogue for Reverse Engineering Assessments and Methods) Challenges focused on reproductive health. We assessed LLM performance across four predictive tasks derived from three DREAM challenges: gestational age regression from gene expression, gestational age regression from DNA methylation profiles, and classification of preterm birth and early preterm birth from microbiome data. LLMs were prompted with task descriptions, data locations, and target outcomes. LLM-generated code was then run to fit and apply prediction models and generate graphics, and they were ranked based on their success in completing the tasks and achieving strong test set performance. Among the eight LLMs tested, o3-mini-high, 4o, DeepseekR1 and Gemini 2.0 completed at least one task without error. Overall, R code generation was more successful (14/16 tasks) than Python (7/16), attributed to the utility of Bioconductor packages for querying Gene Expression Omnibus data. OpenAI’s o3-mini-high outperformed others, completing 7/8 tasks. Test set performance of the top LLM matched or exceeded top-performing teams from the original DREAM challenges. These findings underscore the potential of LLMs to enhance exploratory analysis and democratize access to predictive modeling in omics by automating key components of analysis pipelines, and highlight the potential to increase research output when conducting analyses of standardized datasets from public repositories.

## Introduction

Reproductive health is a critical area of study that encompasses fertility, pregnancy, and childbirth, all of which have profound implications for individual and public health. Among these concerns, preterm birth—defined as delivery before 37 weeks of gestation—remains a major global challenge, affecting approximately 11% of infants worldwide and leading to significant short- and long-term health consequences^1^. Understanding and addressing issues in reproductive health is essential for developing effective prevention strategies, promoting maternal and infant well-being, and reducing health disparities. In recent years, predictive models in reproductive medicine have been developed to estimate the likelihood of adverse pregnancy outcomes, supporting personalized care and informed decision-making^2^. However, the performance of these models depends heavily on the quality and size of the datasets used, the distribution and types of data available, and the specific methods applied^3^. Continued research in reproductive health, supported by robust data and innovative technology, is key to improving outcomes across the reproductive lifespan.

Recent advances in maternal health research underscore the critical importance of accurately estimating gestational age^4^ and understanding the molecular mechanisms underlying placental aging^5^. Precise determination of gestational age is essential for optimizing clinical management, guiding timely interventions, and reducing adverse outcomes for both mother and child. Equally, insights into the “placental clock”—the epigenetic and molecular processes that regulate placental aging—provide invaluable clues to fetal development, maternal adaptation, and long-term health trajectories^6^. Crowdsourced open science initiatives, exemplified by challenges such as the DREAM (Dialogue for Reverse Engineering Assessments and Methods) Challenges^7^, harness global collaboration and diverse datasets, including transcriptomic^8^, microbiome^9^, and methylation data^10^ to develop predictive models addressing these critical questions. While such collaborative approaches offer access to a broad spectrum of data, computational methodologies and collective expertise, they also face challenges related to coding variability and inconsistent model performance.

Large Language Models (LLMs) are a promising solution to issues of inconsistency. Cutting-edge systems such as GPT-4, ChatGPT, and others, have evolved rapidly, and their applications find critical utility in biological and medical sciences where the challenge of “too much data, too few experts” has long hindered progress^11,12^. By utilizing vast corpora of text-based data—spanning scientific literature, experimental results, and patient records—LLMs are enabling breakthroughs in cancer diagnosis, radiology, electronic health record analysis, hypothesis generation, and more [PathChat^13^,Flamingo-CXR^14^, Med-PaLM^15^]. Another application of LLMs is to democratize data analysis to non-expert programmers, streamline analysis and accelerate the discovery and interpretation process in biomedical data sciences. This has been followed by widespread adoption of LLMs in all parts of the scientific process.

Alongside this exciting explosion in LLMs usage comes the need for careful and meticulous evaluation, and many researchers have shown both their potential for breakthrough discoveries, as well as their potential pitfalls^16^.

Herein, we utilized the wealth of crowdsourced benchmark data from DREAM Challenges to assess the ability of LLMs to support the development of code to leverage predictive models with omics data, and then assess and visualize model performance. Each DREAM challenge consisted of a research question coupled with training and test data. Teams from around the world submitted solutions to tackle a problem based on training data, which were then evaluated on a blinded test set, mirroring the train - test process in machine learning. We prompted various popular LLMs to mirror a typical workflow of a bioinformatician who writes R or Python code to read data from local files or web-based repositories, to build and evaluate a prediction model for multiple tasks. The data types considered included transcriptomic, epigenetic, and microbiome, spanning several orders of magnitude in terms of problem dimensionality. We systematically assessed the LLMs performance in a single-shot setting using 4 prediction tasks: Gestational age prediction from blood gene expression or placenta methylation data, and preterm or early preterm birth classification from microbiome data. Executing and evaluating the code generated by LLMs, we derived insight in terms of coding language-specific reliability of the LLM-generated code and their ability to match the test set prediction accuracy of participants in the original DREAM challenges. Our study reframes the usage of LLMs in the context of crowdsourced research efforts, highlighting their ability to generate reproducible machine learning pipelines that reduce inconsistencies and improve efficiency in a multi-team collaborative setting.

## Methods

The methodology used in this study is depicted in Figure 1. Briefly, each of the 8 LLMs considered (Table S1) was prompted to generate R or Python code that uses one training dataset that (i) fits a model for the given task, (ii) applies the model to the corresponding test set to calculate an appropriate performance metric, and (iii) generates a visualization of the result (Table 1). These prompts specified the type and source of data, dimensionality of the feature space, prediction outcome, data partitioning into training and test sets, and required model evaluation metrics [root mean square error (RMSE) for continuous outcomes and area under the curve of the receiver operating characteristics curve (AUC-ROC) for binary classification]. The LLM-generated code was saved and edited by a) adding extra lines of code to enable saving raw predictions and b) disabling code lines attempting to install R or python packages. The code was executed in Python or R, using the same computing system (Red Hat Enterprise Linux 8.10 Ootpa, with 56 logical processors, 250 GiB of RAM) with all R/Python packages required by any LLM already installed. The test set accuracy and generated plots were then scored to enable LLM ranking. The scores were determined by assigning a) 1 point for successful data extraction and formatting, b) 2 points for successful model training and application of the model on the test set and metric calculation, c) 1 point for successful generation of the required plot, d) 1 point for achieving highest prediction accuracy (within ±0.02 for AUC-ROC and ±0.1 for RMSE of the top model for that task).

**Table 1:** Prompts and computational tasks. Prompts used for each prediction task and LLM. For each task, the main goal/endpoint is noted, along with the number of features in the three different datasets used. The exact prompt given to the LLMs is shown.

| Task/Data type | Prompt to LLM |
| --- | --- |
| Q1:<br>Predict gestational age from gene expression in blood<br>~30k features | Write a program in **R/python** which downloads the Gene Expression Omnibus dataset GSE149440 and uses expression data of tens of thousands of genes in a few hundred samples to fit a prediction model of gestational age (gestational age:ch1) variable from the metadata. Only samples assigned to the training set (metadata variable train:ch1=='1') should be used for training. Apply the model on the test defined as metadata variable train:ch1==0. Print the root mean squared error (RMSE) on the test set and generate a scatter plot of the predicted vs actual gestational age values for the test set. |
| Q2<br>Predict gestational age from placenta methylation data<br>~360k features | Write **R/python** code that fits one predictive model for a continuous variable gestational age (GA) using about 400 thousand methylation features measured in about 2000 samples. The feature data for the training set is in df_training.csv while for the test set is in df_test.csv. Features are rows and samples are columns in the feature data, with the first column being the feature names. The metadata files are ano_training.csv and ano_test.csv. The feature data and metadata are linked by the Sample_ID. Do not use any other metadata variables in the models besides feature data. Print the root mean squared error (RMSE) on the test set and generate a scatter plot of the predicted vs actual gestational age values for the test set and include RMSE in the legend. |
| Q3A:<br>Predict Preterm Birth from microbial abundance<br>Q3B:<br>Predict Early Preterm Birth from microbial abundance | Write **R/python** code that fits one predictive model for a binary outcome PTB using about two thousands microbiome features measured in a few hundreds of samples. The feature data for the training set is in training_species_abundance.csv while for the test set is in validation_species_abundance.csv. Discard features not in common between training and validation sets and do not use any other metadata in the models besides microbial feature data. The metadata for the training set is in training_metadata.csv while for the test set in validation_metadata.csv. The feature data and metadata are linked by the specimen column in these datasets. There are multiple observations (specimen) for each subject |
| ~2k features | identified by the column participant_id. Do use only the entry with highest collect_wk value < 32 for each participant_id to fit and evaluate the models. PTB is defined as metadata variable delivery_wk < 37. Print AUC ROC value on the test set and plot an ROC curve. Repeat the analysis above to predict EarlyPTB outcome defined as delivery_wk < 32 using data for last specimen with collect_wk<28 for each participant_id. |

**Figure 1.**
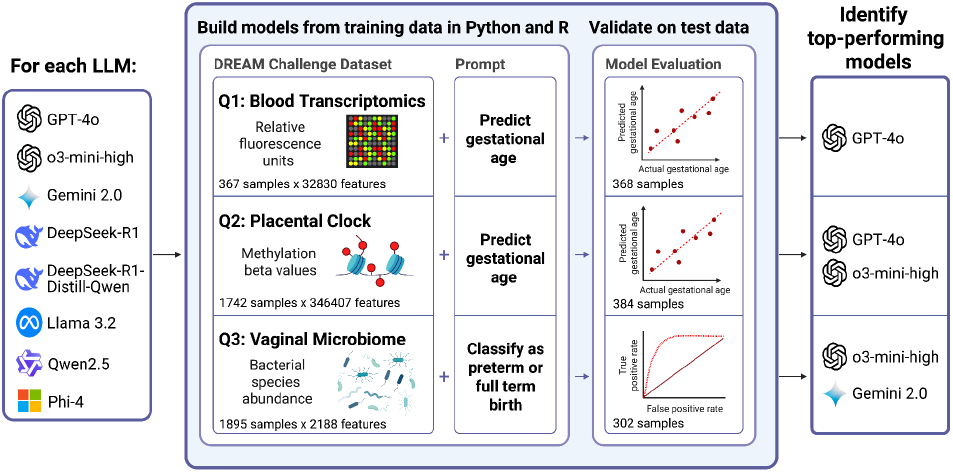
Study flow chart. Eight LLMs were prompted to generate R and Python code to fit and evaluate predictive models for four tasks. Q1: predict gestational age from blood transcriptomics; Q2: predict placenta age from DNA methylation data; Q3A: predict preterm birth or Q3B: early preterm birth from microbial relative abundance data; Analysis code was executed and LMMs were ranked by code functionality and model accuracy. Created with BioRender.

## Results

The prediction tasks used to assess LLMs were based on datasets from multiple biomedical domains, each representing distinct analytical challenges (Figure 1, Table S2). The first dataset (Q1) involved transcriptomics data obtained from the Gene Expression Omnibus (GEO), requiring parsing and preprocessing before downstream analysis. The second dataset (Q2) focused on epigenetics, specifically methylation data for the prediction of placental gestational age using regression models. The third dataset (Q3) was used for two classification tasks based on relative microbial abundance. Each of these prediction tasks tested the ability of LLMs to retrieve and organize data, foresee the need for data conversion, identify a suitable modeling framework, and select the appropriate package for fitting those modeling strategies across different programming environments.

The test set accuracy metrics obtained by executing the code generated by LLMs for all tasks and programming languages are shown in Figure 2. The empty cells in the table signify that the code generated by the specific LLM resulted in an error, preventing the completion of the task. Overall, among the four LLMs that generated code which successfully completed any task (o3-mini-high, 4o, DeepSeekR1, and Gemini), the R code generated by the LLMs was more successful in completing the four tasks (14/16) compared to Python (7/16). Three LLMs created R code that successfully generated models and test set predictions, but failed to produce code to plot results. Plots successfully generated by LLMs for each task are shown in Figures S1-3. Due to the lack of support for parsing the GEO data (Q1) in Python, none of the LLMs were able to complete this task. However, due to the efficient implementations of high-dimensional models (such as Ridge Regression), the RMSE in Python for task Q2 obtained by 4o and DeepSeekR1 was lower than that of the best-performing team in the Placental Clock DREAM Challenge. These results align with previous studies highlighting the effectiveness of high-dimensional regression models in biomedical applications^17^. The best LLM prediction metrics for the other three tasks were worse than those obtained by participants in these DREAM challenges (Q1 RMSE = 5.42 vs. 4.5; Q3A AUROC= 0.57 vs. 0.6; Q3B AUROC= 0.59 vs. 0.6) (Figure 3). Out of all the LLMs we tested, o3-mini-high received the highest overall score (33), followed by 4o (23), DeepSeekR1 (22), and Gemini2.0 FlashThinking (19) (Table 2).

**Table 2.**
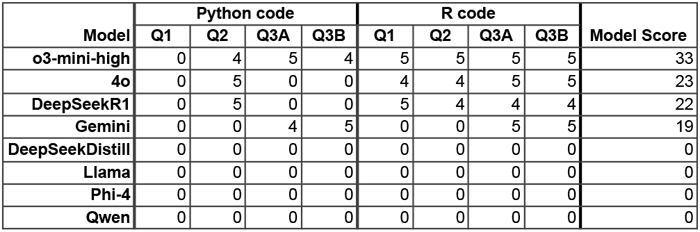
Model scoring. For each task (Q1, Q2, Q3A, Q3B), models were evaluated for their ability to produce Python and R code that successfully extracted and formatted data, model training, application to the test set, performance metric calculation, generation of the required plot, and prediction accuracy.

| Model | Python code |  |  |  | R code |  |  |  | Model Score |
| --- | --- | --- | --- | --- | --- | --- | --- | --- | --- |
|  | Q1 | Q2 | Q3A | Q3B | Q1 | Q2 | Q3A | Q3B |  |
| **o3-mini-high** | 0 | 4 | 5 | 4 | 5 | 5 | 5 | 5 | 33 |
| **4o** | 0 | 5 | 0 | 0 | 4 | 4 | 5 | 5 | 23 |
| **DeepSeekR1** | 0 | 5 | 0 | 0 | 5 | 4 | 4 | 4 | 22 |
| **Gemini** | 0 | 0 | 4 | 5 | 0 | 0 | 5 | 5 | 19 |
| **DeepSeekDistill** | 0 | 0 | 0 | 0 | 0 | 0 | 0 | 0 | 0 |
| **Llama** | 0 | 0 | 0 | 0 | 0 | 0 | 0 | 0 | 0 |
| **Phi-4** | 0 | 0 | 0 | 0 | 0 | 0 | 0 | 0 | 0 |
| **Qwen** | 0 | 0 | 0 | 0 | 0 | 0 | 0 | 0 | 0 |

**Figure 2.**
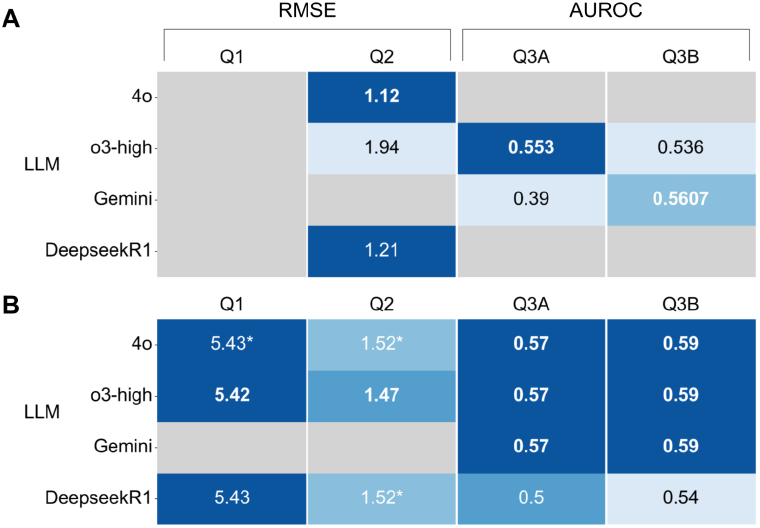
Test set prediction results. Accuracy of models generated by executing LLM-generated Python (A) and R (B) analysis code. Colors are normalized by dataset; best scores are indicated by the darkest color per column, and the best score per dataset and language is shown in bold. Gray cells indicate that the code generated an error, preventing the completion of the task. Asterisks indicate that the model did not successfully plot the results. Models listed in Table S1 but not included here did not generate any successful code.

**Figure 3.**
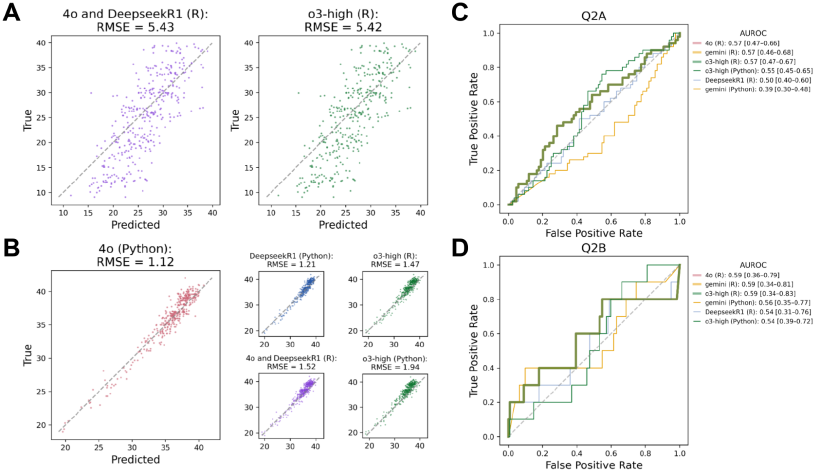
AUROC and RMSE curves. A. Predicted vs. actual gestational ages for all successful LLMs for Q1. B. Predicted vs. actual placental gestational ages for the top scoring LLM for Q2 (left) and lower scoring LLMs (right subplots). 4o- and DeepseekR1-generated code output identical predictions for A and B and are plotted together in purple. C. ROC curves for all LLM-generated Q3A predictions. D. ROC curves for all LLM-generated Q3B predictions. In C and D, the three highest scoring models output identical predictions and are represented by the bold green line. Area Under the Receiver Operating Characteristic curve (AUROC) [95% confidence interval (CI) lower bound, upper bound] for each model was calculated via 1,000 bootstrap resamples of the true and predicted labels. Confidence intervals were estimated by resampling the prediction dataset with replacement and computing AUCs on each sample. All model CIs show substantial overlap, indicating comparable performance across models.

## Discussion

Our benchmarking study leveraging -omics data from DREAM Challenges highlights both the promise and limitations of LLMs in biomedical machine learning applications. Similar benchmarking approaches have been employed to assess LLM performance in other domains, including chemistry and clinical medicine^18,19^. Herein, we found that API-based LLMs such as o3-mini-high and 4o outperformed smaller locally-run models in execution success and accuracy, reinforcing the advantages of larger, cloud-hosted architectures. However, local models offer increased control and customization, which may be advantageous for researchers seeking data privacy or fine-tuned execution. Prior research has shown similar trade-offs in LLM performance across different computing environments^20^.

In terms of test set prediction accuracy of models obtained with LLM-generated analysis code compared to human-generated code, our results suggest that in some instances, LMM-generated models can be more accurate and faster to develop. For task Q2, where the LLM-generated model from ∼350k methylation features was more accurate than the human counterpart (RMSE=1.12 vs 1.24, respectively), neither the LLM nor the human had access to the test set or had any feedback on their model accuracy based on the test set. For the other three prediction or classification tasks, the human participants had the advantage of either using additional feature data in the models (Q3A and B), or receiving and using feedback on their model accuracy based on the test set (Q1 and Q3A and B).

One of the key motivations for integrating LLMs into the biomedical research process is their ability to support computational tasks by allowing prototyping of analysis code for researchers without coding skills and improving the productivity of those with coding skills. While crowdsourcing has democratized access to datasets and allows assessment of models on blinded data while facilitating global collaboration, it also introduces challenges related to reproducibility, coding variability, and quality control^21^. LLMs offer a potential solution by generating standardized, executable code, reducing the variability inherent in manually coded solutions. Furthermore, LLMs can automate complex preprocessing steps, ensuring consistency in data handling across multiple modeling approaches, a known limitation of crowdsourced competitions^22^.

One of the possible limitations of the use of LLMs in the context of data challenges such as DREAM, is that multiple identical models may be obtained from different LLMs by different users, if the prompts are the same or similar. This was illustrated in this study for task Q1 where 3 models achieved the same accuracy (RMSE 5.4 weeks). Multiple identical models will lead to ties in the ranking of the participating teams and less diversity in the approaches and ultimately less insight into the question formulated in the DREAM challenge. However, the use of LLMs by scientists in their daily analysis tasks has the potential to streamline model development and reduce over-fitting, particularly in reproductive health research, where accurate and reproducible predictive models can significantly impact clinical decision-making, and reliable biomarkers derived from omics studies are scarce. Future research should explore how LLMs can be further fine-tuned for biomedical applications, particularly in the context of preterm birth and maternal-fetal medicine. Additionally, the integration of LLM-generated models into clinical pipelines should be rigorously evaluated to assess their translational potential and ensure alignment with medical best practices.

This study provides a comprehensive benchmarking of LLMs for biomedical predictive modeling, evaluating their ability to generate machine learning code across multiple data modalities and coding languages. By addressing key challenges associated with code and models generated from crowdsourcing in biomedical research, LLMs offer a promising avenue for improving reproducibility, standardization, and coding efficiency, yet they may lead to identical models generated by challenge participants who adopt LLMs. While API-based LLMs demonstrated superior reliability, locally run models provide opportunities for enhanced control. LLMs offer significant potential in automating machine learning workflows but require careful validation to ensure reproducibility and accuracy in biomedical research. Ongoing advancements in LLM architectures and prompt engineering strategies are poised to further refine their utility in scientific computing, including efforts to address major public health challenges such as preterm birth.

## Code availability

https://github.com/vtarca7/LLMDream

## Data availability

See Table S2

