## Supplemental Table and Figures for "Benchmarking Large Language Models for Predictive Modeling in Biomedical Research With a Focus on Reproductive Health"

**Supplementary Tables**

**Table S1: LLMs used in the comparison**

| LLM | Developer | Parameters | Run Method |
|---|---|---|---|
| Deepseek-R1 | DeepSeek | 671B | API |
| 4o | OpenAI | Unknown | API |
| o3-mini-high | OpenAI | Unknown | API |
| Gemini2.0 FlashExpThink | Google | Unknown | API |
| Qwen2.5 Coder | Alibaba | 14B | Local |
| Llama3.2 | Meta | 3B | Local |
| Phi-4 | Microsoft | 14B | Local |
| DeepSeek-R1-Distill-Qwen | DeepSeek | 32B | Local |

**Table S1:** Details the eight different LLMs tested for generating code for the 8 different prediction tasks. Developers of each LLM are listed, as well as the method by which the LLM was run (API vs. Local). When available, information on the number of parameters for each LLM is described as well.

**Table S2: Dataset info**

| Dataset | DREAM Challenge name | Accession Number | Number of samples | | |
|---|---|---|---|---|---|
| | | | Train | Test | Total |
| **Q1** | Predict gestational age from blood transcriptomics data | SDY2187 GSE149440 | 367 | 368 | 735 |
| **Q2** | Predict gestational age from placental methylation data | SDY2187 | 1742 | 384 | 2126 |

| | | | | | |
|---|---|---|---|---|---|
| **Q3 (A and B)** | Classify term vs. preterm birth from vaginal microbiome data | SDY2187 SDY465 | 1895 | 302 | 2197 |

**Table S2:** Details the four different tasks each LLM was prompted to perform. Q1 task prompted LLMs to build a model to predict gestational age from transcriptomics data, Q2 prompted LLMs to predict gestational age again, but from methylation data, and Q3 prompted LLMs to build a model to classify preterm birth from microbiome data. The table also details the study accession numbers for each of the datasets, as well as the number of samples in the training, test, and full datasets. SYD refers to synapse identifier (synapse.org), while GSE is a Gene Expression Omnibus identifier.

### Supplementary Figures

**A** **B**

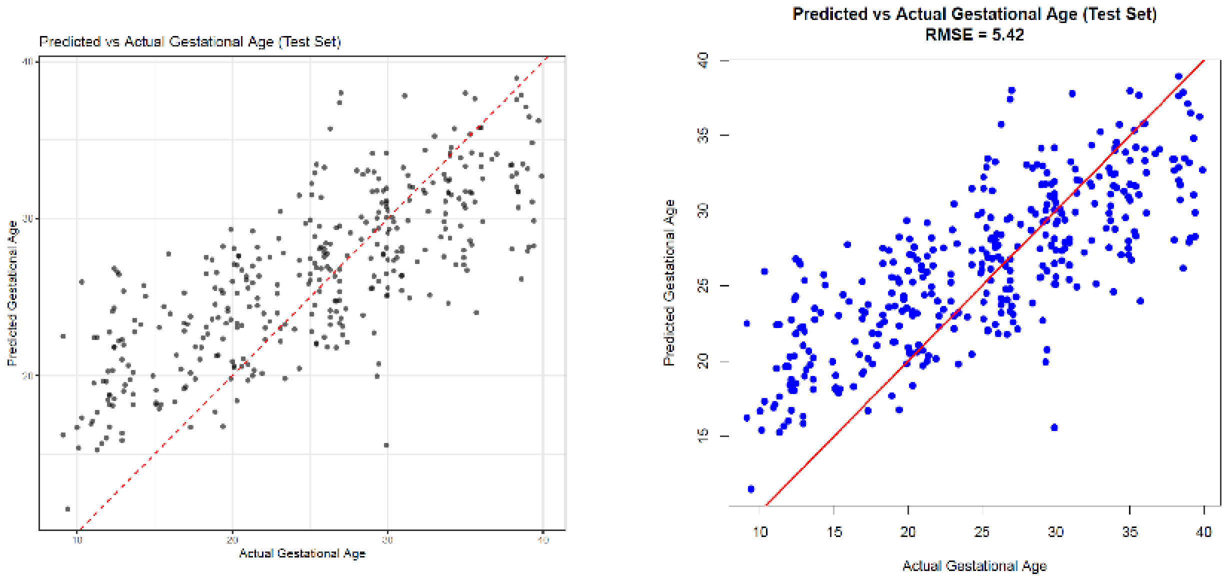


**Figure S1:** (A) Predicted vs. actual gestational age from blood transcriptomics data, produced by DeepseekR1, run in R. RSME was 5.43. (B) Predicted vs. actual gestational age produced by GPT o3-mini-high model, run in R. RSME of 5.42.

**A** **B**

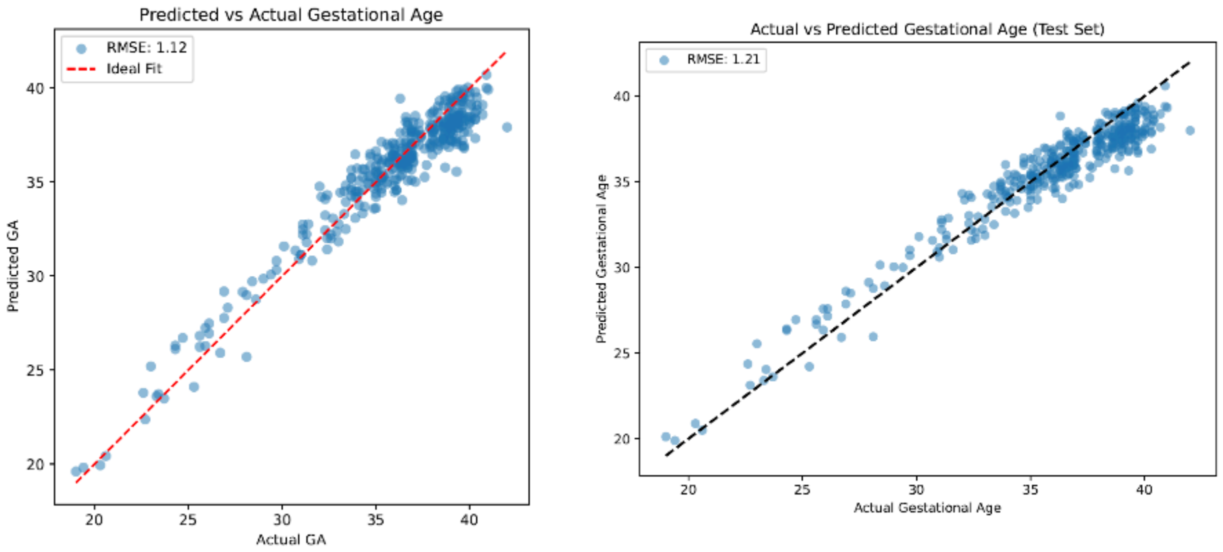


**Figure S2:** A. Predicted vs. actual gestational age from placental methylation data. (A) GPT 4o model results, run in Python. RSME of 1.12. (B) DeepseekR1, run in Python. RSME of 1.21. Both models had better performance than the original top crowdsource-produced model (RMSE=1.24).

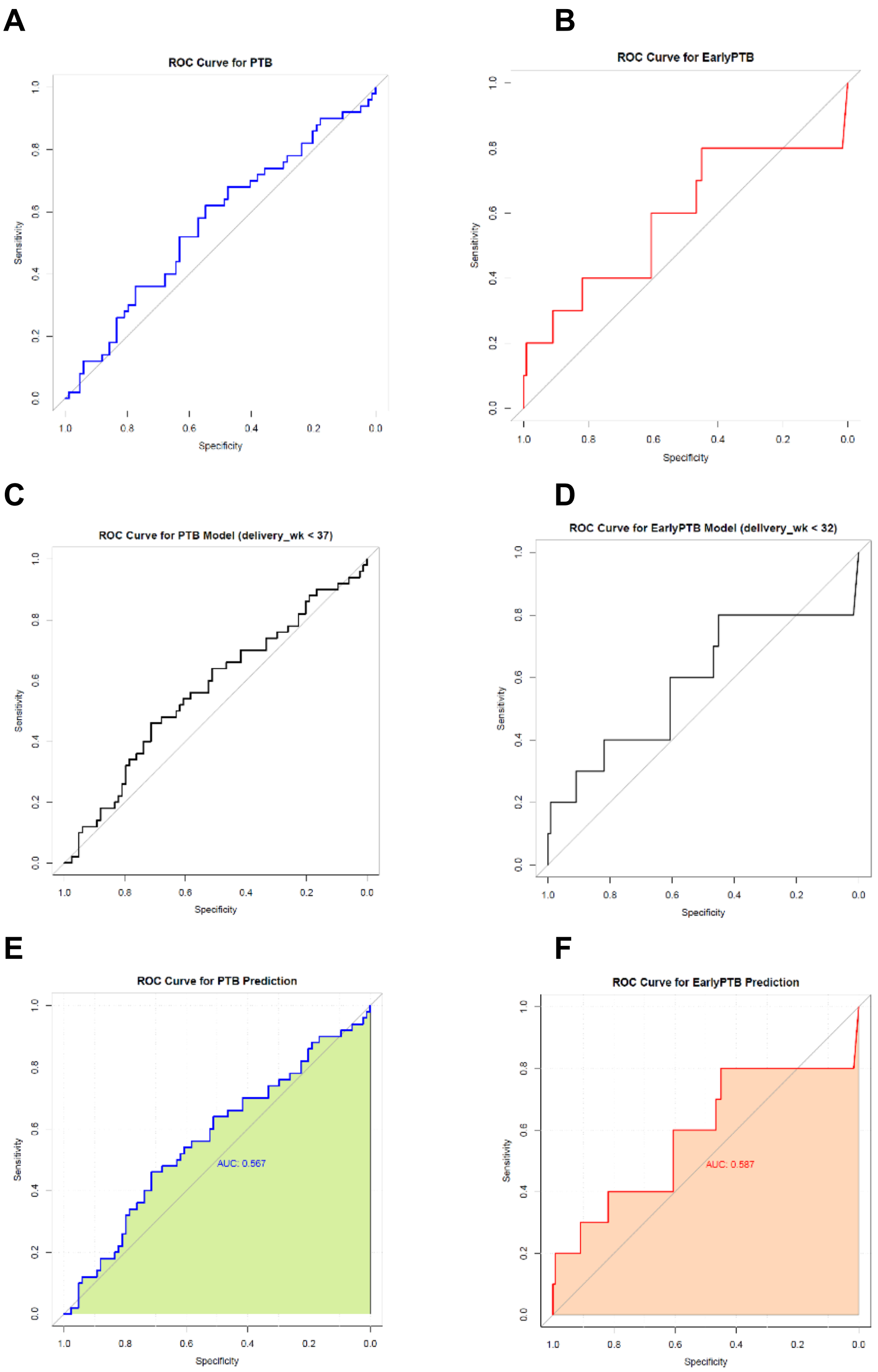


**Figure S3:** ROC curves for classification of preterm birth and early preterm birth produced by different LLMs, where all code was run in R. (A,B). AUC ROC curve produced from GPT-4o's model for (A) preterm birth classification and (B) early preterm birth classification. (C,D). AUC ROC curve produced from o3-mini-high model for (C) preterm birth classification and (D) early preterm birth classification. (E,F). ROC curve produced from Gemini’s model for (E) preterm birth classification and (F) early preterm birth classification.
